# Impact of gene annotation choice on the quantification of RNA-seq data

**DOI:** 10.1101/2021.01.07.425794

**Authors:** David Chisanga, Yang Liao, Wei Shi

## Abstract

RNA sequencing is currently the method of choice for genome-wide profiling of gene expression. A popular approach to quantify expression levels of genes from RNA-seq data is to map reads to a reference genome and then count mapped reads to each gene. Gene annotation data, which include chromosomal coordinates of exons for tens of thousands of genes, are required for this quantification process. There are several major sources of gene annotations that can be used for quantification, such as Ensembl and RefSeq databases. However, there is very little understanding of the effect that the choice of annotation has on the accuracy of gene expression quantification in an RNA-seq analysis. In this paper, we present results from our comparison of Ensembl and RefSeq human annotations on their impact on gene expression quantification using a benchmark RNA-seq dataset generated by the SEquencing Quality Control (SEQC) consortium. We show that the use of RefSeq gene annotation models led to better quantification accuracy, based on the correlation with ground truths including expression data from >800 real-time PCR validated genes, known titration ratios of gene expression and microarray expression data. We also found that the recent expansion of the RefSeq annotation has led to a decrease in its annotation accuracy. Finally, we demonstrated that the RNA-seq quantification differences observed between different annotations were not affected by the use of different normalization methods.

## 1 INTRODUCTION

Gene expression profiling using RNA sequencing (RNA-seq) is a core activity in molecular biology. Comprehensive gene expression analysis in various settings is important for generating hypotheses for ongoing research, investigating drug-effects in biological or clinical settings and as a diagnostic tool. In this paper, we explore the fact that a popular approach in gene-level quantification from RNA-seq data involves mapping reads to a reference genome and then counting mapped reads associated with each gene [1, 2, 3, 4, 5]. The process of counting mapped reads to genes requires a database of known genes. A gene is only quantified if it or its components have genomic coordinates already defined with respect to the genome sequence in a process called annotation. For each genome annotation model, a different set of annotation techniques and information sources are used and as such, these annotations vary in terms of comprehensiveness and accuracy of annotated genomic features. Annotation techniques often include computer-based predictions and/or evidence-based techniques such as manual curation [6, 7]. Computer-based predictions result in more complex gene models that have a higher proportion of predictive genomic features while evidence-based generated gene models are simpler with fewer genes and isoforms. Common annotation models for human and mouse genomes include Ensembl [8], RefSeq [9], GENCODE [10] and UCSC [11] annotations. Annotations are, therefore, an important component in an RNA-seq analysis as the results are dependent on what is known in the annotation database.

Despite the importance of gene annotations in RNA-seq data analysis, very little research has been conducted to examine how differences in annotations impact on gene expression quantification, which is crucial for downstream analyses such as discovery of differentially expressed genes and identification of perturbed pathways. Previous studies compared the effect of human genome annotations from popular databases including Ensembl, GENCODE and RefSeq on various aspects of RNA-seq analysis and they showed that the choice of annotations had an impact on gene-level quantification in the RNA-seq analysis [12, 13]. However, these studies are out of date as they were based on old annotations and they also lacked a reliable ground truth for assessing the impact of annotation.

Major annotation databases have undergone significant expansions over the years, thanks to the wide application of sequencing technologies and the massive amount of sequencing data that have been generated across the world. However, it is unclear whether the quality of gene annotations have been successfully maintained. A recent study suggested that gene annotations have become less accurate and lagging during this expansion [6]. This can be attributed to the errors from sequencing experiments, sequence analysis or automation in the annotation process. It is important to systematically assess the accuracy of the new gene annotations generated in recent years to ensure the popular annotation databases can continue to be utilized by the community for RNA-seq analysis.

Furthermore, the use of different annotations in different studies makes it difficult for researchers to reproduce the findings from such studies. For example, large consortia such as the European Molecular Biology Laboratory (EMBL) use Ensembl in their studies while the National Centre for Biotechnology Information (NCBI) tend to use RefSeq. Since this can significantly impact on gene expression data, there is a need to develop a comprehensive understanding of how these differences in annotations impact the gene-level expression quantification.

In this study, we compared three human gene annotations, including a recent Ensembl annotation (released in April 2020), a recent RefSeq annotation (released in August 2020) and an old RefSeq annotation (released in April 2015), to understand their impact on gene-level expression quantification in an RNA-seq data analysis pipeline. Although the old RefSeq annotation is not available at the NCBI RefSeq database anymore, it has been included as part of Rsubread, a popular RNA-seq quantification toolkit, for quantifying human RNA-seq data. We used a benchmark RNA-seq dataset generated by the SEquencing Quality Control (SEQC/MAQC III) consortium for this evaluation. We show that the use of RefSeq gene annotations led to better quantification accuracy than the use of Ensembl annotation, based on the correlation with ground truths including expression data from >800 real-time PCR validated genes, known genome-wide titration ratios of gene expression and microarray gene expression data. We also show that the older RefSeq annotation yielded higher quantification accuracy than the recent RefSeq annotation in our evaluations, suggesting that the recent expansion and changes made to the RefSeq annotation have led to a decline in annotation accuracy resulting in less accurate quantification result. Furthermore, we investigated if any normalization method can mitigate the differences in quantification results caused by the annotation differences. Our results show that the quantification differences remained almost the same no matter how the RNA-seq data were normalized.

## 2 MATERIALS AND METHODS

### 2.1 SEQC/MAQC data

The RNA-seq data used for evaluation in this study are a benchmark dataset generated by the Sequencing Quality Control (SEQC) project [1], the third stage of the MicroArray Quality Control (MAQC) study [14, 15]. The SEQC dataset includes the Universal Human Reference RNA (UHRR) as sample A and the Human Brain Reference RNA (HBRR) as sample B. It also includes two other samples C and D, which are combination of A and B mixed in the ratios of 3:1 in C and 1:3 in D respectively. The samples were sequenced in four replicate paired-end libraries using an Illumina HiSeq 2000 sequencer at the Australian Genomics Research Facility (AGRF). Each library contains ~20 million 100bp read pairs.

A TaqMan real-time polymerase chain reaction (RT-PCR) dataset with expression values measured for over 1,000 genes, which was generated in the MAQC-I study [15], was used to validate the expression of the RNA-seq data in this study. The expression values were measured for both the UHRR and HBRR samples together with their respective combinations. Around 800–900 TaqMan RT-PCR genes, which had matching gene identifiers with expressed RNA-seq genes from different annotations, were included for assessing the accuracy of RNA-seq quantification. In addition, microarray data generated in the MAQC-I study with samples A to D hybridized to the Illumina Human-6 Bead-Chip microarrays were also used in the assessment. The TaqMan RT-PCR and Illumina microarray datasets are available as part of the Bioconductor package ‘seqc’ [16].

### 2.2 Annotations used

Three human gene annotations were included in this study, including a recent Ensembl annotation, a recent RefSeq annotation and an old RefSeq annotation. All these annotations were generated based on the human reference genome GRCh38/hg38.

The Ensembl gene annotation used in this study was generated in April 2020. Its version number is 100. It was downloaded from ftp://ftp.ensembl.org/pub/release-100/gtf/homo_sapiens/Homo_sapiens.GRCh38.100.gtf.gz.

The recent RefSeq gene annotation used was released by the NCBI in August 2020. Its release number is 109.20200815 and it is part of the RefSeq release version 202. It was downloaded from the NCBI FTP site ftp://ftp.ncbi.nlm.nih.gov/refseq/H_sapiens/annotation/annotation_releases/109.20200815/GCF_000001405.39_GRCh38.p13/GCF_000001405.39_GRCh38.p13_genomic.gtf.gz. We refer this RefSeq annotation as ‘RefSeq-NCBI’ in this study.

The old RefSeq annotation included in this study was released by the NCBI in April 2015. It was released as part of the Patch 2 release of the GRCh38/hg38 genome build. This annotation has also been included in the popular RNA-seq quantification toolkit Rsubread [5] as the default annotation used for quantifying human RNA-seq data. The inclusion of this old RefSeq annotation allowed us to investigate how the annotation changes made recently to RefSeq affect the quantification result of RNA-seq data. The RefSeq annotation in Rsubread is slightly different from the original one in that the overlapping exons from the same gene were collapsed to form a single continuous exon for the gene in the Rsubread annotation, however this difference will not change the gene-level RNA-seq quantification result because the set of exonic bases belonging to each gene is the same between the original annotation and the Rsubread annotation. As this old RefSeq annotation is no longer available for downloading at the NCBI FTP site, we instead used the Rsubread annotation in this study and we denote this annotation as ‘RefSeq-Rsubread’.

When matching genes from different annotations, we converted the gene identifiers using the Bioconductor package ‘org.Hs.eg.db’ [17] and then compared them to find common genes between annotations.

### 2.3 Mapping, quantification and normalization of RNA-seq data

Analysis of the RNA-seq data was performed using Bioconductor R packages Rsubread and limma [5, 18, 19]. The human reference genome (GRCh38) from GENCODE (version 34 downloaded from ftp://ftp.ebi.ac.uk/pub/databases/gencode/Gencode_human/release_34/GRCh38.primary_assembly.genome.fa.gz) was indexed using the buildin-dex function in Rsubread v2.2.6 [5]. Sequencing reads were then mapped to the reference genome using the align function in Rsubread [5, 20]. During the alignment, the Ensembl, RefSeq-NCBI and RefSeq-Rsubread annotations were also included as an extra parameter to improve alignment.

Gene-level read counts were obtained with featureCounts [4, 5], a read count summarization function within the Rsubread package. The Ensembl, RefSeq-NCBI and RefSeq-Rsubread annotations were provided to featureCounts to generate read counts for genes included in these annotations respectively.

The gene-level read counts were transformed using the voom function in limma [18, 21] and then normalized using the library size [22], quantile [23] and trimmed mean of M-values (TMM) [24] methods, respectively, prior to performing further analysis. The library size normalization was performed by providing raw read counts to voom and then running voom with the ‘normalize.method’ parameter set to ‘none’. The quantile normalization was performed by providing raw read counts to voom and then running voom with the ‘normalize.method’ parameter set to ‘quantile’. For TMM normalization, we first calculated the TMM normalization factor for each library using the calcNormFactors method in edgeR [25]. Then we provided raw read counts and the TMM normalization factors to voom and ran it with the ‘normalize.method’ parameter set to ‘none’. The log_2_CPM (log_2_ counts per million) values, produced by the voom function for each gene in each library, were converted to log_2_FPKM (log_2_ fragments per kilo exonic bases per million mapped fragments) expression values for further analysis.

### 2.4 Titration monotonicity

The RNA-seq data from the SEQC project have titration monotonicity built into them, such that a gene is considered to preserve titration monotonicity if the expression of the gene follows A ≥ C≥ D ≥B when its expression in sample A is greater than or equal to that in sample B, or follows A ≤ C≤ D ≤B when its expression in sample A is less than or equal to that in sample B. To test if the titration monotonicity is preserved, Equation (1) was used to compute the expected log2 fold-change for a gene in the comparison of C vs D given the log2 fold-change between A vs B.

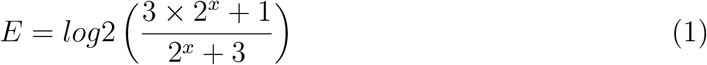

where *E* is the expected log2 fold-change for C vs D and *x* is the log2 fold-change for A vs B. Expression levels of genes in the replicates of the same sample were averaged before fold change of gene expression was calculated between samples.

### 2.5 Validation

Gene expression data generated using TaqMan RT-PCR and Illumina’s BeadChip microarray were used to validate the gene-level quantification results from the RNA-seq analysis. Pearson correlation coefficients were computed to assess the concordance between the RNA-seq quantification data obtained from using different annotations and the gene expression data obtained from the RT-PCR and microarray experiments. The genome-wide built-in truth of titration monotonicity of gene expression in the RNA-seq data was also utilized to evaluate the quantification accuracy of RNA-seq data generated from using different annotations.

### 2.6 Access to data and code

The data and analysis code used in this study can be accessed at the following URL:

https://github.com/ShiLab-Bioinformatics/GeneAnnotation.

## 3 RESULTS

### 3.1 Discrepancy between different gene annotations

The Ensembl and NCBI RefSeq annotations are among the most widely used gene annotations that have been utilized for RNA-seq gene expression quantification in the field. In this study, we downloaded recent Ensembl and RefSeq annotations and also used an older version of Refseq annotation to assess the impact of gene annotation choice on the accuracy of RNA-seq expression quantification. The inclusion of an older RefSeq annotation allowed us to investigate the accuracy of new annotation data generated in recent years when the next-gen sequencing data have been used as a new data source for genome-wide annotation generation.

The Ensembl annotation used in this study was released in April 2020 and it has a version number 100. The recent RefSeq annotation included in this study was released in August 2020. We call this annotation as ‘RefSeq-NCBI’ in this study. The older RefSeq annotation was released in April 2015, and it has also been included as part of the popular RNA-seq quantification toolkit ‘Rsubread’ for quantifying human RNA-seq data. As this annotation is not available in the NCBI RefSeq database anymore, we instead used the Rsubread RefSeq annotation in our evaluations and we denote this annotation as ‘RefSeq-Rsubread’.

As RNA-seq gene-level expression quantification is typically performed for genes that contain exons [3, 4, 5], in this study we only focused on the genes that have annotated exons in each annotation. Figure 1A shows that, as expected, the Ensembl annotation contains a lot more exon-containing genes than the two RefSeq annotations. The Ensembl annotation is known to contain a large number of computationally predicted genes whereas RefSeq genes were mainly annotated based on the biological evidence. However, it is worth noting that the RefSeq-NCBI annotation still has >14,000 genes that are not included in the Ensembl annotation. Nearly 60% of the Ensembl genes were found to be absent from both of the two RefSeq annotations. In total, 23,424 common genes were found between the three annotations. Most of the genes included in the RefSeq-Rsubread annotation can be found in the RefSeq-NCBI or Ensembl annotations.

**Figure 1:**
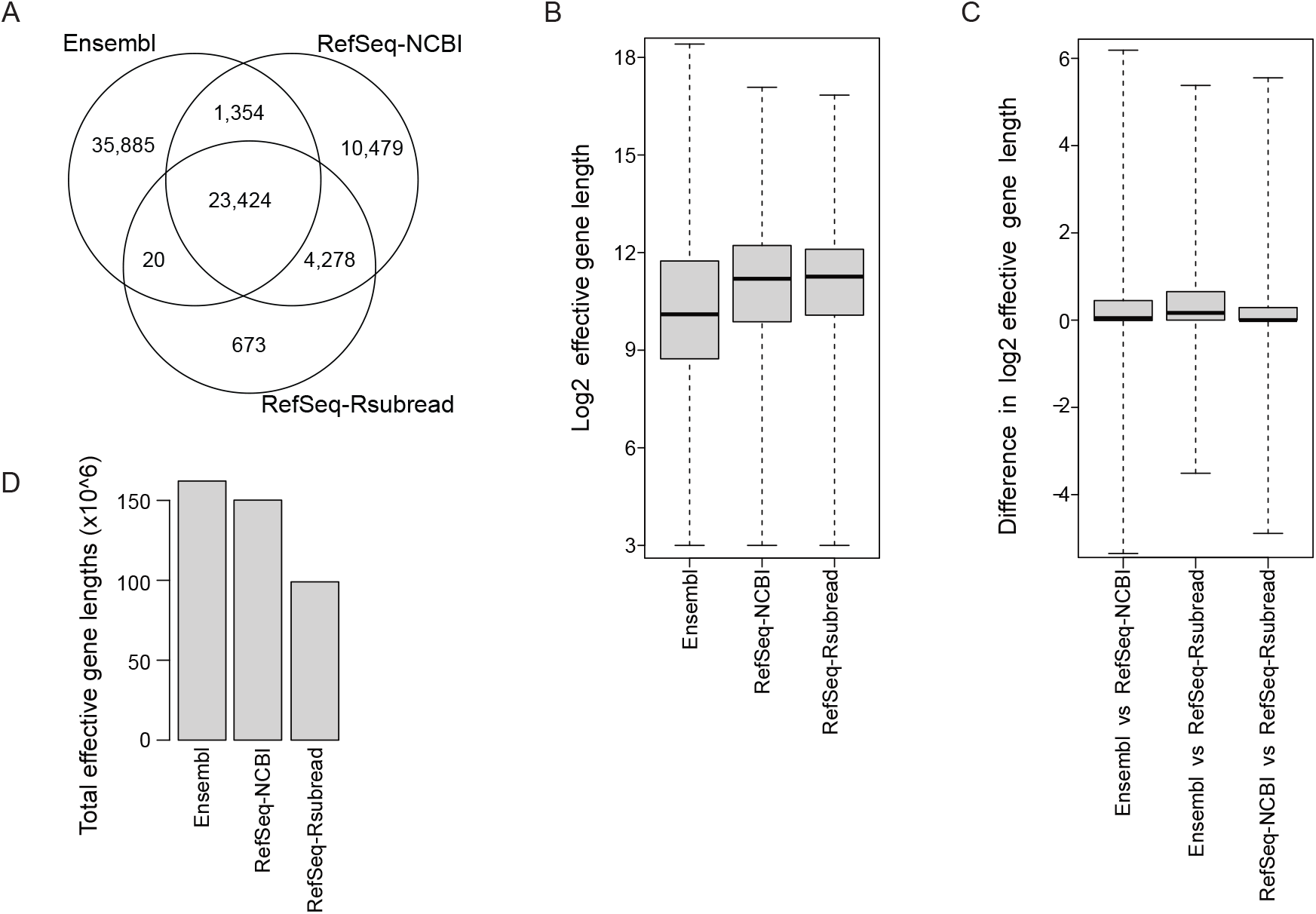
Concordance and differences between gene annotations. (A) Venn diagram showing genes that are common or unique in the Ensembl, RefSeq-NCBI and RefSeq-Rsubread annotations. (B) Boxplots showing the distribution of effective gene lengths (*log*_2_ scale) in each annotation. (C) Boxplots showing the differences in effective lengths of common genes between each pair of annotations. Values shown in the plots are the ratio of effective lengths of the same gene from two different annotations (*log*_2_ scale). (D) The size of transcriptome calculated from each annotation. Shown are the sum of effective gene lengths in each annotation.

We then examined the effective gene lengths in each annotation. The effective length of a gene is the total number of unique bases included in all the exons belonging to the gene. Figure 1B shows the distributions of effective lengths of genes in the three annotations. Around half of the Ensembl genes have an effective length less than 1,000 bases, whereas in the two RefSeq annotations only ~25% of the genes are shorter than 1,000 bases in length. The median effective gene lengths in RefSeq-NCBI and RefSeq-Rsubread are ~3,000 bases, which is much larger than that in Ensembl (~1,000 bases). Although the Ensembl annotation contains a lot more genes than the two RefSeq annotations, it also contains a much higher percentage of short genes.

We further performed gene-wise comparison of effective gene lengths using common genes between each pair of annotations. Although every annotation contains both longer and shorter genes in comparison to the corresponding genes from other annotations, the Ensembl genes were found to have a larger effective length than genes from the two RefSeq annotations overall (Figure 1C). This is in contrast to the higher proportion of short genes observed in the Ensembl annotation (Figure 1B), which indicates that the Ensembl genes that are also present in RefSeq-NCBI or RefSeq-Rsubread annotations tend to be longer than those Ensembl genes that can only be found in the Ensembl annotation. Although at least half of the genes were found to have a less than 2-fold (1-fold at *log*_2_ scale) length difference between annotations (Figure 1C), the length differences could be as high as more than 64-folds (6-folds at *log*_2_ scale). The RefSeq-NCBI genes seem to be slightly longer than the corresponding RefSeq-Rsubread genes overall. Ensembl and RefSeq-Rsubread were found to be the least concordant annotations among the three annotations being compared.

Lastly, we compared the size of the transcriptome represented by each annotation. The transcriptome size of an annotation is computed as the sum of effective gene lengths from all the genes included in that annotation, which also represents the total number of exonic bases that were annotated in an annotation. Figure 1D shows that the Ensembl annotation has a larger transcriptome size than both RefSeq-NCBI and RefSeq-Rsubread annotations. This is not surprising because the Ensembl annotation contains more genes and also Ensembl genes common to other annotations are longer in general. RefSeq-Rsubread has a much smaller transcriptome size than RefSeq-NCBI, indicating a significant expansion of the RefSeq-NCBI annotation in the past five years. However, it is important to note that the RefSeq-Rsubread annotation is not a subset of the RefSeq-NCBI annotation, as demonstrated by the existence of RefSeq-RSubread genes that are absent in the RefSeq-NCBI annotation, the difference in gene length distribution and the length differences of the same genes between the two annotations (Figure 1A-C). This indicates that not only were new genes added to the RefSeq annotation during the expansion, but existing genes have been modified.

It is against this background that we sought to understand how these differences in the annotations impact on the overall gene-level quantification results.

### 3.2 Fragments counted to annotated genes

We used a benchmark RNA-seq dataset generated by the SEQC project [1] to evaluate the impact of gene annotation on the accuracy of RNA-seq expression quantification. This dataset contains paired-end 100bp read data generated for four samples including a Universal Human Reference RNA sample (sample A), a Human Brain Reference RNA sample (sample B), a mixture sample with 75%A and 25%B (sample C) and a mixture sample with 25%A and 75%B (sample D).

We mapped the RNA-seq reads to the human genome GRCh38/hg38 using the Sub-read aligner [20, 5], and then counted the number of mapped fragments (read pairs) to each gene in each annotation using the featureCounts program [4, 5]. FeatureCounts assigns a mapped fragment to a gene if the fragment overlaps any of the exons in the gene. Figure 2 shows that across all the 16 libraries, the RefSeq-Rsubread annotation constantly has substantially more fragments assigned to it than the Ensembl and RefSeq-NCBI annotations. This is surprising because RefSeq-Rsubread contains much less annotated genes and also has a significantly smaller transcriptome, compared to Ensembl and RefSeq-NCBI (Figure 1A,D). We then performed a detailed investigation into the mapping and counting results to find out what enabled RefSeq-Rsubread to achieve a higher percentage of successfully assigned fragments.

**Figure 2:**
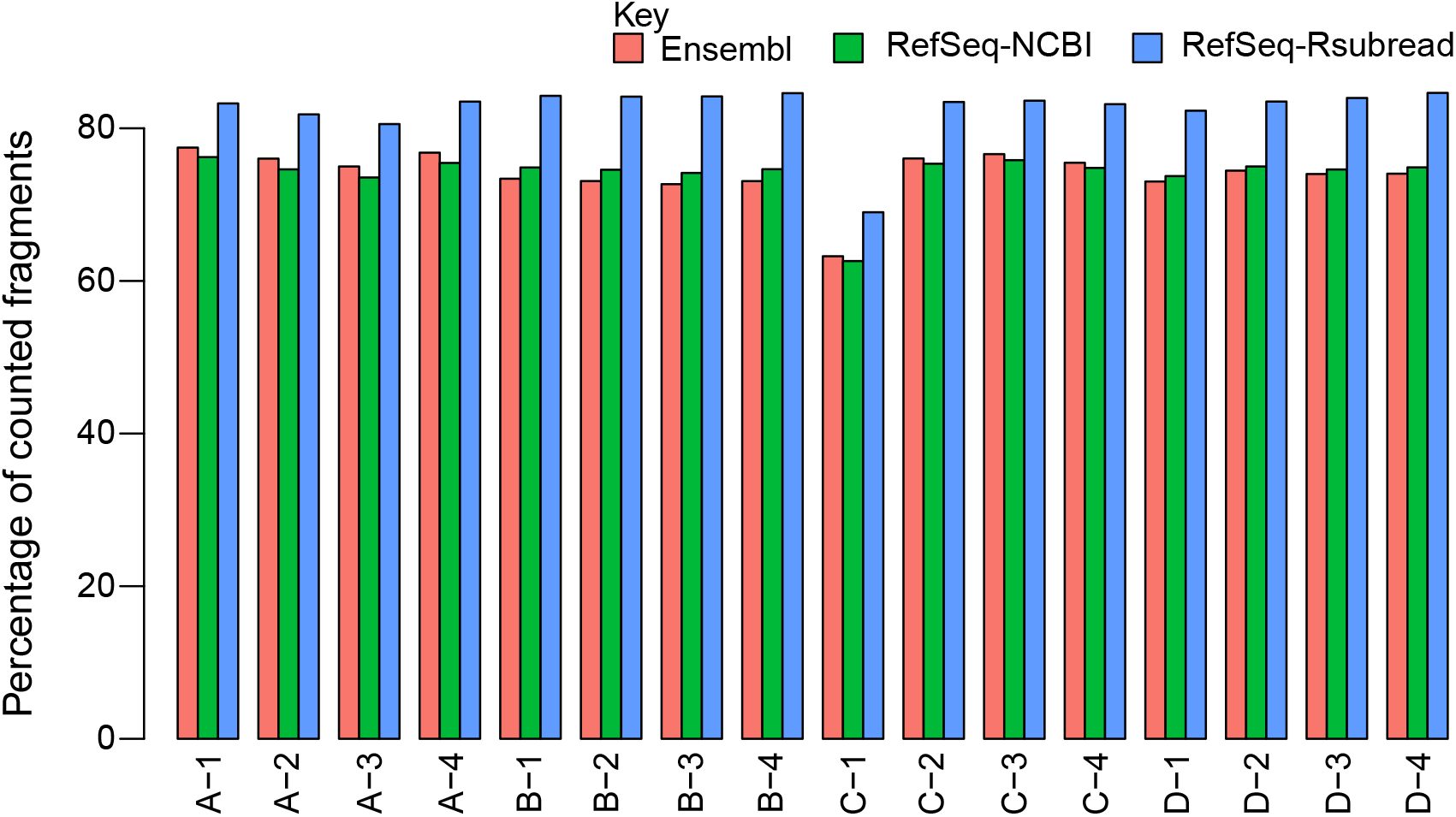
Barplots showing the percentage of fragments successfully assigned to genes in each annotation, out of all the fragments included in each library. The horizontal axis represents the sixteen SEQC RNA-seq libraries generated from the four samples ‘A’, ‘B’, ‘C’ and ‘D’. Each sample has four replicates that are numbered from 1 to 4.

Although gene annotations were utilized in mapping reads to the human reference genome, the use of different annotations was not found to affect the number of successfully aligned fragments for each library (Supplementary Figure S1). We found that when assigning fragments to genes using the Ensembl or RefSeq-NCBI annotation, more fragments were unable to be assigned because they did not overlap any genes (ie. failed to overlap any exons included in any genes), despite there are more genes included in these annotations compared to the RefSeq-Rsubread annotation (Supplementary Figure S2). This is particularly the case for the fragment assignment in the human brain reference samples. We also found that the use of Ensembl and RefSeq-NCBI annotations led to more fragments being unassigned due to the assignment ambiguity, ie. a fragment overlaps more than one gene (Supplementary Figure S3). This should be because there are more genes that overlap with each other (ie. exons from different genes overlap with each other) in the Ensembl and RefSeq-NCBI annotations compared to the RefSeq-Rsubread annotation. Our investigation revealed that less gene overlapping in the RefSeq-Rsubread annotation and better compatibility of this annotation with the mapped fragments have led to more fragments being successfully counted for each library in this dataset. Given that both the Universal Human Reference and Human Brain Reference samples used in this study are known to contain a very high number of expressed genes and the RNA-seq data generated from these samples are expected to cover most of the human transcriptome, our analysis suggests that the RefSeq-Rsubread annotation is likely to contain more transcribed region in the genome than the other two annotations in general.

### 3.3 Intensity range of gene expression

We examined if the gene annotation choice has an impact on the range of gene expression levels in the RNA-seq data. Raw gene counts of the SEQC data were converted to log_2_FPKM (log_2_ fragments per kilo exonic bases per million mapped fragments) values for all the genes included in each annotation. A prior count of 0.5 was added to the raw counts to avoid log-transformation of zero. Figure 3 shows that the two RefSeq annotations exhibit a desirable larger intensity range of gene expression than the Ensembl annotation, as shown by the larger boxes in the boxplots. It is surprising to see that the Ensembl genes have the smallest intensity ranges in all the libraries, give that the Ensembl annotation contains the largest number of genes in all the three annotations being examined. In addition to the large intensity range, the RefSeq-Rsubread genes were also found to have a markedly higher median expression level than genes in the RefSeq-NCBI and Ensembl annotations.

**Figure 3:**
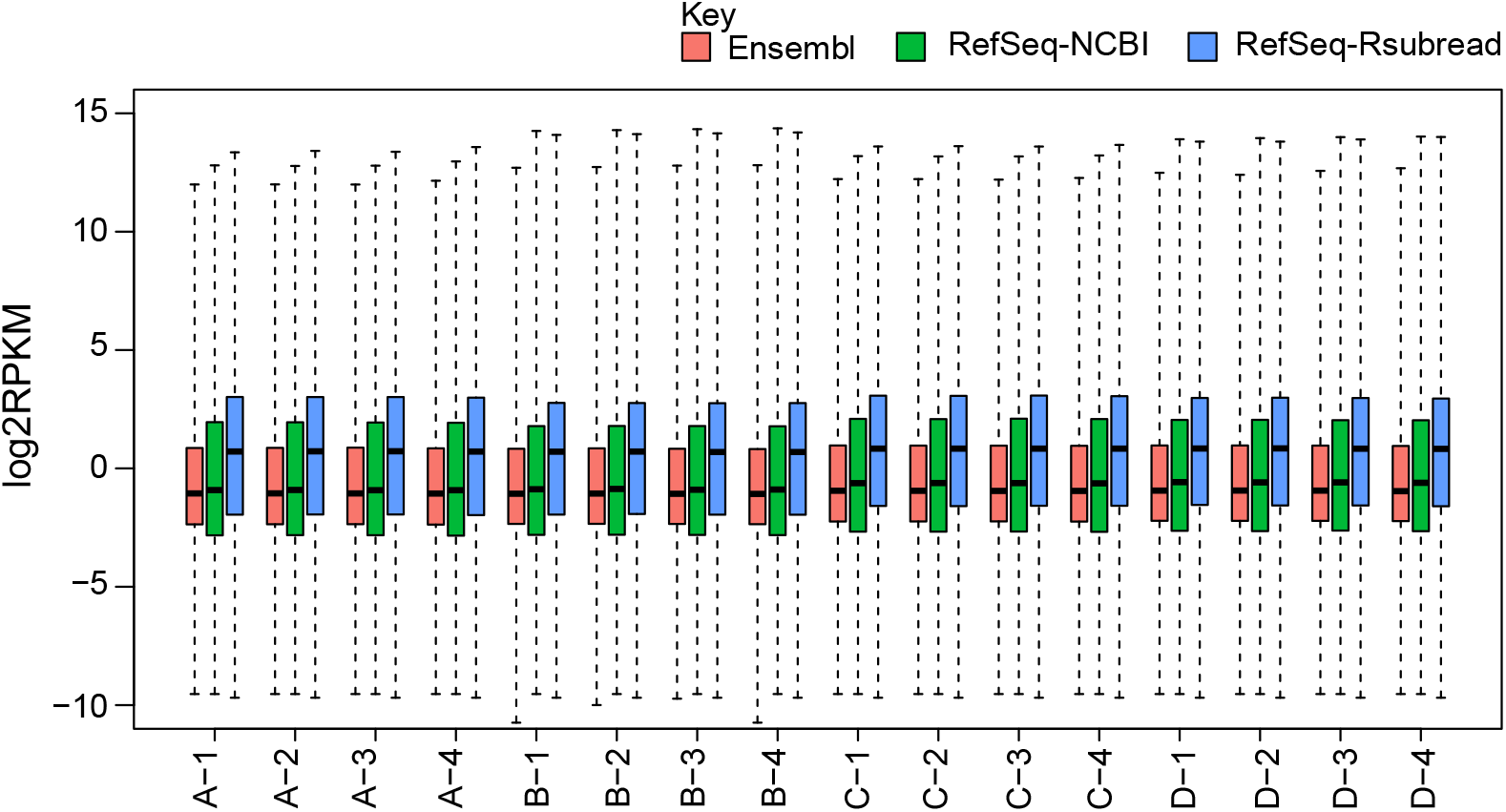
Boxplots comparing the intensity range of gene expression between the three annotations. All the genes from each annotation were included in the plots. Raw read counts of genes were transformed to log_2_FPKM values. A prior count of 0.5 was added to raw counts to avoid log-transformation of zero.

### 3.4 Gene annotation discrepancy after expression filtering

As it is a common practice to filter out genes that are deemed as lowly expressed, or are completely absent in an RNA-seq data analysis [2], we also set out to assess the differences between alternative annotations after excluding such genes. We excluded those genes that failed to have at least 0.5 CPM (counts per million) in at least four libraries (each sample has four replicates) in the analysis of the SEQC dataset. The expression-filtered data were also used for comparing the accuracy of quantification from using alternative annotations presented in the following sections.

The bar plot in Figure 4A shows that Ensembl has significantly more genes (also higher proportion of genes) filtered out due to low or no expression, compared to RefSeq-NCBI and RefSeq-Rsubread. After expression filtering, the total numbers of remaining genes from the three annotations became more similar to each other. 16,472 genes were found to be common between the three annotations after filtering, accounting for 69%, 78% and 86% of the filtered genes in the Ensembl, RefSeq-NCBI and RefSeq-Rsubread annotations respectively (Figure 4B). Almost all the filtered genes in the RefSeq-Rsubread annotation can be found in the other two annotations.

**Figure 4:**
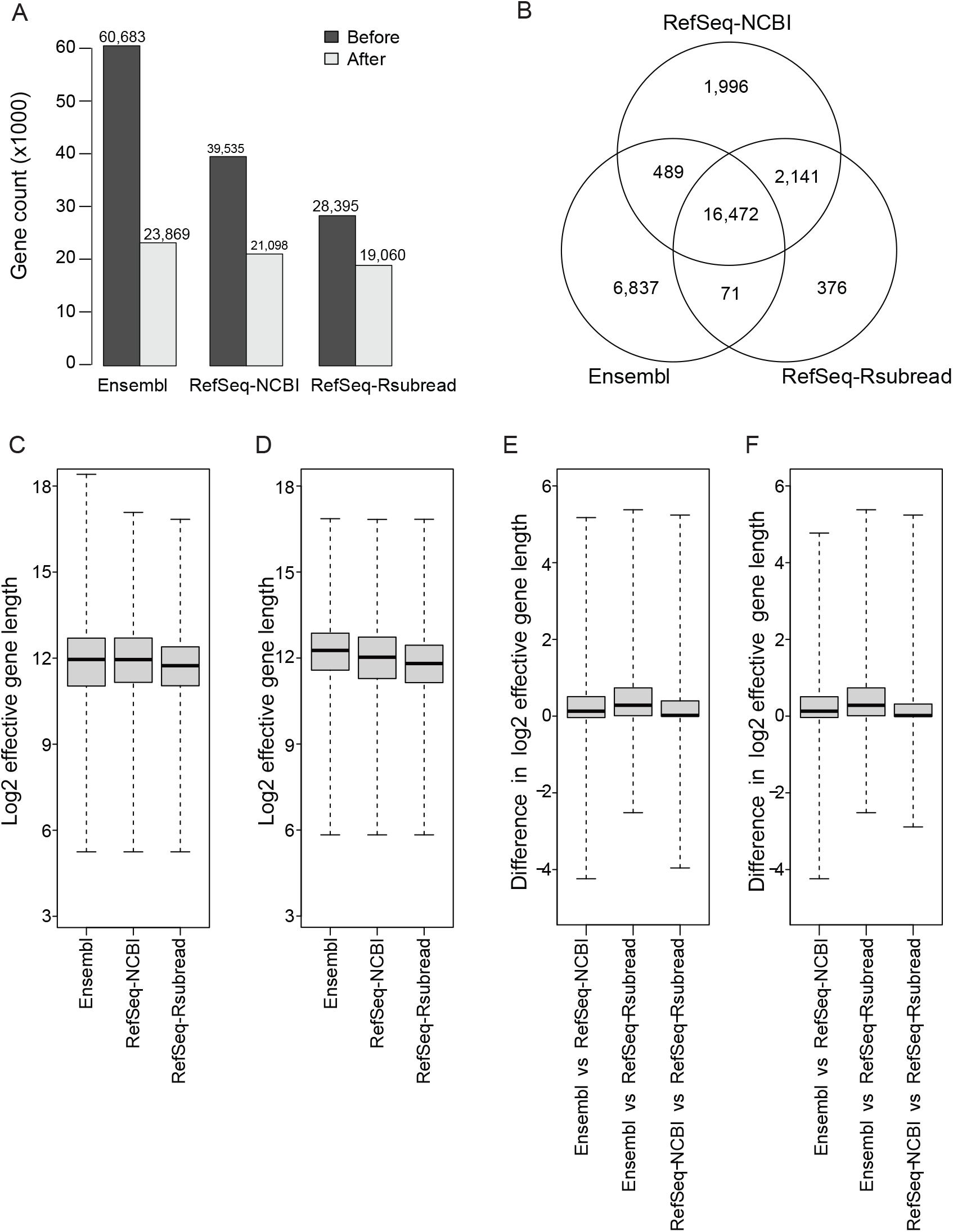
Concordance and differences between gene annotations after filtering for lowly expressed genes. (A) Bar plot showing the differences in the number of genes included in each annotation before and after filtering for lowly expressed genes. (B) Venn diagram comparing genes from different annotations after filtering for lowly expressed genes. Distributions of effective gene lengths after filtering are shown for all genes in each annotation (C) and for genes that are common between all three annotations (D). Distributions of differences of effective gene lengths between annotations after filtering are shown for common genes between each pair of annotations (E) and for genes that are common between all three annotations (F).

After expression filtering, the median effective gene length has increased to ~4,000 bases for all annotations (Figure 4C), meaning that a higher proportion of short genes were removed due to low expression in every annotation. The median effective length of Ensembl genes now became comparable to, or slightly higher than those in the two RefSeq annotations, indicating that the Ensembl annotation contained a higher proportion of lowly expressed short genes than the two RefSeq annotations. When comparing the effective lengths of genes common to all three annotations after filtering, the Ensembl genes were found to have the largest median effective length and the RefSeq-Rsubread genes have the smallest median effective length (Figure 4D). This is not surprising because the Ensembl annotation is known to be more aggressive than the RefSeq annotations and the RefSeq-Rsubread annotation is an old annotation that has not been updated in the last five year.

The expression filtering did not seem to affect the distribution of differences of effective gene lengths between each pair of annotations (using genes common to each pair of annotations), with Ensembl and RefSeq-Rsubread remaining to be the least concordant annotations (Figure 4E and Figure 1C). Using genes common to all three annotations after filtering exhibited similar distributions of gene length differences between each pair of annotations compared to using genes common to each pair of annotations (Figure 4F). Similar to before filtering, the gene-wise length comparison performed after filtering also showed that overall the Ensembl genes had the largest gene lengths and the RefSeq-Rsubread genes had the shortest gene lengths.

### 3.5 Comparison of titration monotonicity preservation

To assess the impact of gene annotation choice on the accuracy of RNA-seq quantification result, we utilized as ground truth the inbuilt titration monotonicity in the SEQC data, the TaqMan RT-PCR data and the microarray data generated for the same samples, to evaluate which annotation gives rise to a better expression correlation of the RNA-seq quantification data with the truth.

In this section, we compared the ability of Ensembl and the two RefSeq annotations in retaining the inbuilt titration monotonicity in the RNA-seq dataset. In Figure 5, the reference titration curve depicts the expected fold change that genes are expected to follow in sample C vs sample D based on the fold change in sample A vs sample B. This is computed using the Equation (1) (see MATERIALS AND METHODS). We then calculated the Mean Squared Error (MSE) between the reference titration monotonicity and the titration monotonicity obtained from each annotation. A smaller MSE value means that the generated quantification data is closer to the truth. Figure 5 shows that the MSE computed for the RefSeq-Rsubread annotation is constantly lower than those computed for the Ensembl and RefSeq-NCBI annotations, regardless if filtering was applied or if only common genes were included for comparison. RefSeq-Rsubread was also found to yield comparable or lower MSE compared to the other two annotations when the data were TMM or quantile normalized (Supplementary Figures S4 and S5), in addition to the library-size normalized data shown in Figure 5. These results demonstrated that the use of RefSeq-Rsubread annotation led to better quantification accuracy for the RNA-seq data.

**Figure 5:**
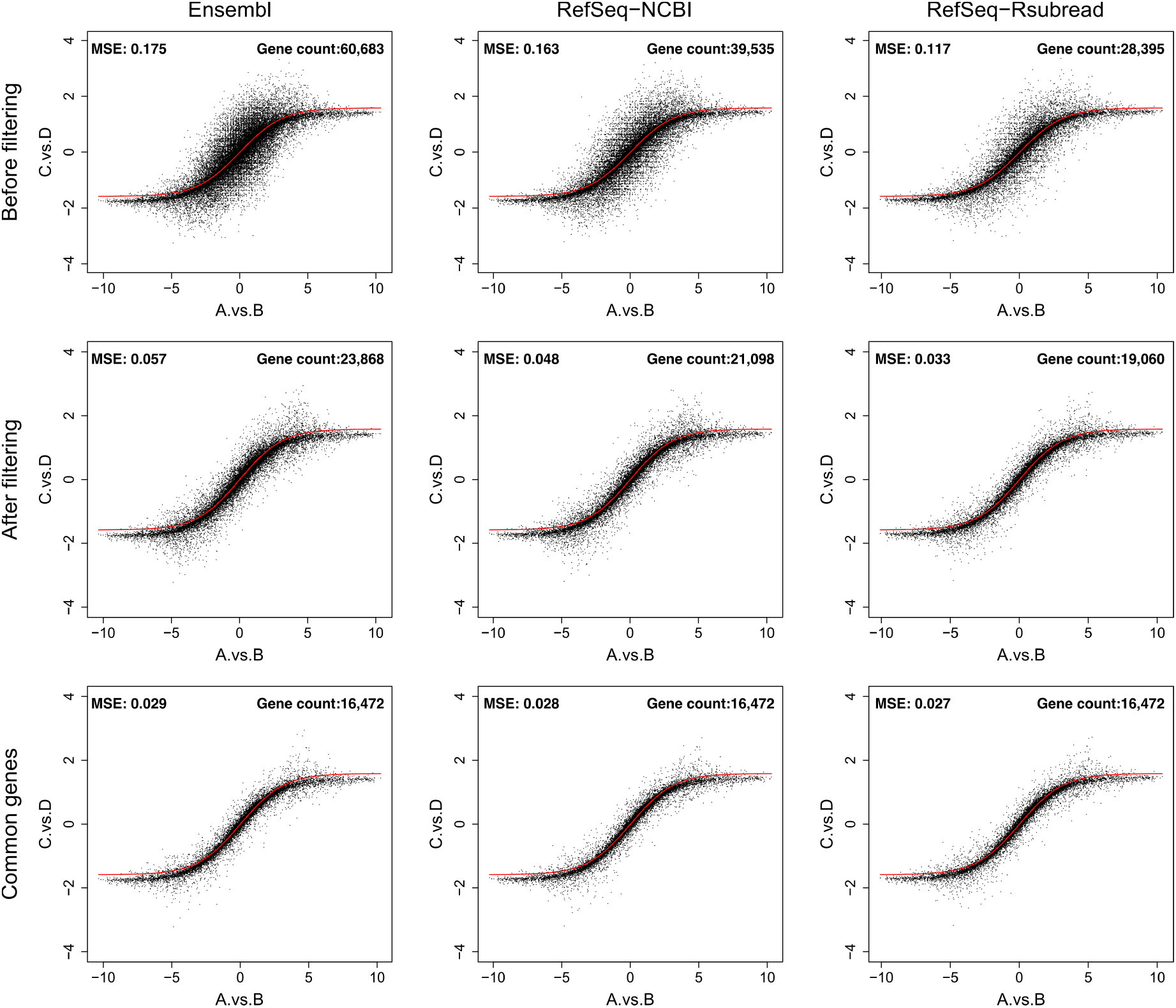
Titration monotonicity plots. The ability of Ensembl, RefSeq-NCBI and RefSeq-Rsubread to retain the titration monotonicity built into the SEQC RNA-seq data was measured using the Mean Squared Error (MSE) between the reference titration and the actual titration obtained from each annotation. The red curve in each plot represents the reference titration calculated from using Equation (1). Plots in the top row include all the genes available in each annotation. Plots in the middle row includes those genes that remained after filtering for lowly expressed genes, in each annotation. Plots in the bottom row includes genes that are common between the three annotations after the expression filtering was performed. In each plot, the horizontal axis represents the log_2_ fold changes of gene expression between samples A and B and the vertical axis represents the log_2_ fold changes of gene expression between samples C and D.

### 3.6 Validation against TaqMan RT-PCR data

The TaqMan RT-PCR dataset generated in the MAQC study [14, 15] was used to validate the gene-level quantification results from the RNA-seq dataset. This dataset contains measured expression levels for >1,000 genes in the four SEQC samples. The aim was to understand how well Ensembl and RefSeq annotated gene expression correlated with the TaqMan RT-PCR data.

The RNA-seq data generated from each annotation were filtered to remove lowly expressed genes before being compared to the RT-PCR data. Numbers of matched genes between the RT-PCR data and the RNA-seq data were 856, 901 and 901 for Ensembl, RefSeq-NCBI and RefSeq-Rsubread, respectively. 846 RT-PCR genes were found to be common to all the three annotations. The raw TaqMan RT-PCR data were log_2_-transformed before comparing to the filtered RNA-seq data.

Pearson correlation analysis of the RNA-seq gene expression (log_2_FPKM values) and RT-PCR gene expression (log_2_ values) from using the RT-PCR genes matched with each individual annotation showed that the RefSeq-Rsubread annotation constantly yielded a higher correlation than the Ensembl and RefSeq-NCBI annotations, across all the samples and the three different normalization methods (left panel in Figure 6). The Ensembl annotation was found to produce the worst correlation in all these comparisons. When using the RT-PCR genes matched with all three annotations for comparison, RefSeq-Rsubread was again found to yield the highest correlation (right panel in Figure 6). Ensembl and RefSeq-NCBI were found to produce similar correlation coefficients. Taken together, results from this evaluation showed that the use of RefSeq-Rsubread annotation led to a better concordance in gene expression between the RNA-seq data and the RT-PCR data, compared to the use of Ensembl and RefSeq-NCBI annotations.

**Figure 6:**
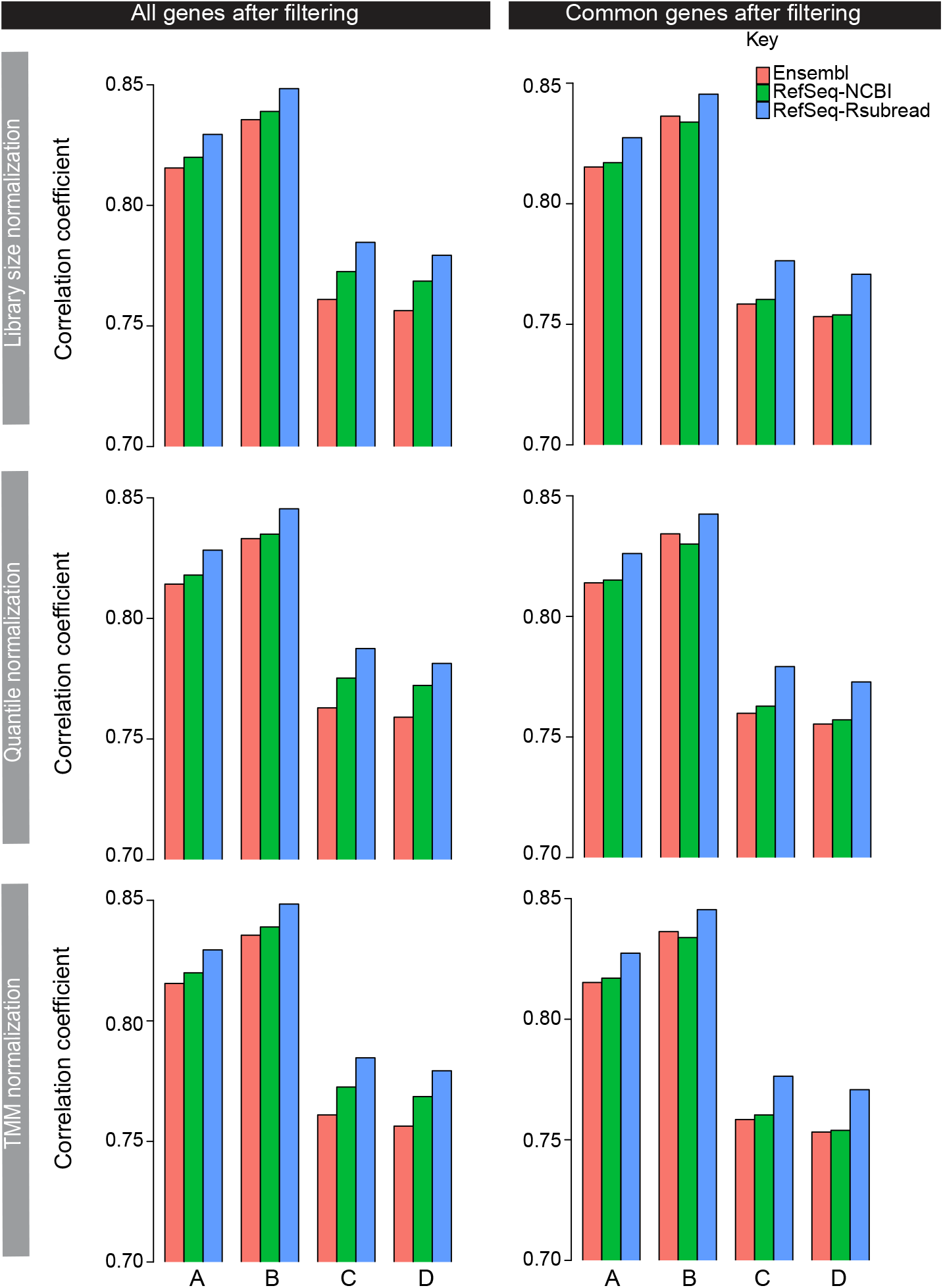
Validation of RNA-seq against TaqMan RT-PCR dataset. Shown are Pearson correlation coefficients computed from comparing RNA-seq data against RT-PCR data, using the RT-PCR genes matched with each individual annotation (left column) or matched with all three annotations (right column). The rows represent the different RNA-seq normalization methods used. Lowly expressed genes in the RNA-seq data were filtered out before the correlation analysis was performed.

### 3.7 Validation against microarray data

An Illumina BeadChip microarray dataset, which was generated by the MAQC-I project [15] for the same samples as in the RNA-seq data used in this study, was used to further validate the gene-level RNA-seq quantification results obtained from different annotations. The microarray dataset was background corrected and normalized using the ‘neqc’ function in the limma package [26, 18]. Microarray genes were then matched to the RNA-seq genes included in the filtered RNA-seq data. 14,405, 14,561 and 14,508 microarray genes were found to be matched with RNA-seq genes from Ensembl, RefSeq-NCBI and RefSeq-Rsubread annotations, respectively. 13,424 microarray genes were found to be present in all three annotations. For those microarray genes that contain more than one probe, a representative probe was selected for each of them. The representative probe selected for a gene had the highest mean expression value across the four samples among all the probes the gene has.

A Pearson correlation analysis was then performed between microarray data and RNA-seq data for each of the three annotations. Both RNA-seq and microarray data include log_2_ expression values of genes. Figure 7 shows that the use of RefSeq-Rsubread annotation consistently yielded the highest correlation between RNA-seq and microarray data in all the comparisons, no matter which RNA-seq normalization method was used and if all or common matched genes were included in the evaluation. On the other hand, the use of the Ensembl annotation resulted in the worst correlation between RNA-seq data and microarray data in all the comparisons.

**Figure 7:**
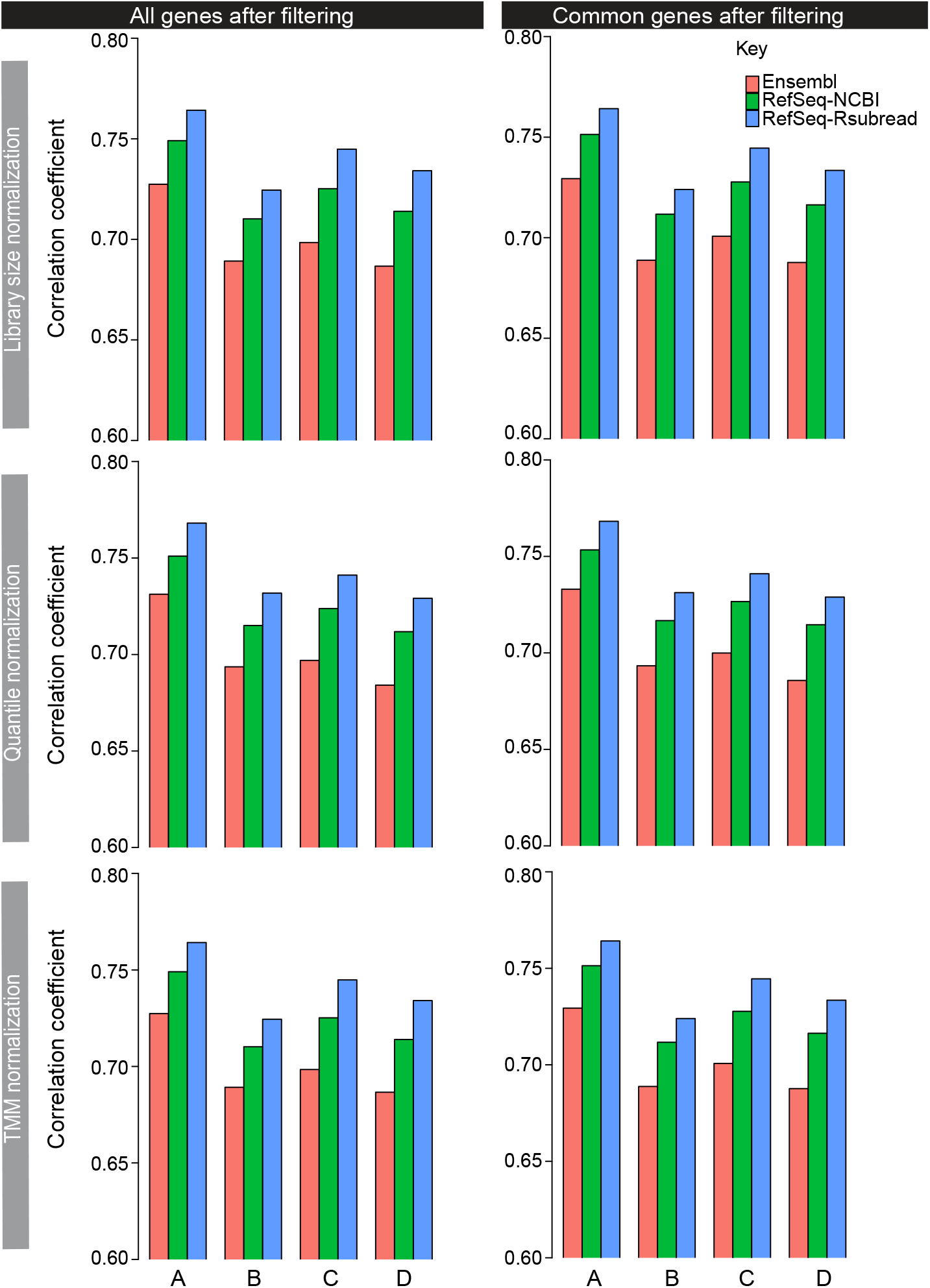
Validation of RNA-seq quantification results against microarray data. Shown are Pearson correlation coefficients computed from comparing RNA-seq data against Illumina BeadChip microarray data, using the microarray genes matched with each individual annotation (left column) or matched with all three annotations (right column). Rows in the plots represent the different RNA-seq normalization methods used. Lowly expressed genes in the RNA-seq data were filtered out before the correlation analysis was performed. For those microarray genes that include more than one probe, a representative probe was selected and used for this analysis.

## 4 DISCUSSION

The RNA-seq technique is currently routinely used for genome-wide profiling of gene expression in the biomedical research field. The analysis of RNA-seq data relies on the accurate annotation of genes so that expression levels of genes can be accurately and reliably quantified. There are several major gene annotation sources that have been widely adopted in the field such as Ensembl and RefSeq annotations. The Ensembl and RefSeq annotations have been well maintained and under continuous development. In particular, new gene information collected from the next-generation sequencing technologies, such as RNA-seq, has been incorporated into the expansion of these annotations in recent years. However, differences between these annotations have raised concerns over the quality and reproducibility of RNA-seq data analyses. There are particularly concerns regarding the accuracy of new gene annotations generated from the use of the sequencing technologies, due to known errors in the generation and analysis of the sequencing data. To address these concerns, in this study we systematically assessed the differences in RNA-seq quantification results attributed to the gene annotation discrepancy. Annotations being evaluated in this study included recent Ensembl and NCBI RefSeq annotations and also an older version of the RefSeq annotation. We compared the recent and old RefSeq annotations to assess the quality of the new annotations that were added when the sequencing technology was utilized at NCBI for curating RefSeq gene annotations.

Although the Ensembl annotation contains significantly more genes than both the recent and old RefSeq annotations, it was also found to have a much higher proportion of short genes. Interestingly, we found that a much higher fraction of these short genes in Ensembl were filtered out due to low or no expression in the analysis of the SEQC RNA-seq dataset, compared to the short genes included in the two RefSeq annotations. The SEQC RNA-seq data is a widely used benchmark dataset including the Human Brain Reference RNA and Universal Human Reference RNA samples, in which a very large number of gene expressed making the entire human transcriptome well covered.

The use of the RefSeq-Rsubread annotation (the older version of the RefSeq annotation used in this study) has led to substantially more fragments being successfully counted to genes than the use of RefSeq-NCBI (the recent RefSeq annotation used in this study) or Ensembl annotations. A detailed investigation revealed that this was because (a) there are less overlapping between genes in the RefSeq-Rsubread annotation leading to less read assignment ambiguity and (b) the RefSeq-Rsubread annotation contains more genes that are compatible with mapped fragments, despite the transcriptome represented by this annotation is much smaller than those represented by the RefSeq-NCBI and Ensembl annotations. Moreover, the quantification data obtained from using RefSeq-Rsubread exhibited desirable larger intensity range and higher median expression level than the quantification data obtained from using the other two annotations.

The evaluation of quantification accuracy from using genome-wide titration monotonicity truth built in the RNA-seq data, the TaqMan RT-PCR data and the microarray data, showed that overall the RefSeq-NCBI annotation yielded better quantification results than the Ensembl annotation. This may not be surprising because the NCBI RefSeq annotation is a traditionally conservative annotation that is known to be highly accurate as it uses an evidence-based approach to annotate genes. However, we also found that the RefSeq-Rsubread annotation yielded more accurate quantification results than the RefSeq-NCBI annotation in almost all the comparisons, which is very surprising. We suspect that this might be due to the annotation errors arising from the sequencing data recently utilized in the NCBI RefSeq annotation generation pipeline. It was reported that the sequencing data, including RNA-seq data and epigenome sequencing data, started to be utilized by NCBI for curating RefSeq gene annotations in around 2013 [9, 27]. Between March 2015 and July 2020, the number of gene transcripts in the vertebrate mammalian organisms included in the RefSeq database increased significantly from 3.6 million to 7.8 million (https://www.ncbi.nlm.nih.gov/refseq/statistics/), a more than twofold increase in just around 5 years. The use of sequencing data for annotation generation should be a significant driver for this rapid expansion of the RefSeq database. It is known that some errors associated with the generation and analysis of sequencing data are difficult to correct, such as sample contamination, sequencing errors, read mapping errors and read assembly errors. When these errors were brought to the annotation process, they could result in incorrect gene annotations being generated and consequently led to less accurate quantification of the RNA-seq data.

## 5 CONCLUSION

In conclusion, our findings from this study revealed that the NCBI RefSeq human gene annotations outperformed the Ensembl human gene annotation in the quantification of RNA-seq data. However, we also raised concerns over the recent changes made to the RefSeq database due to the use of sequencing data in the annotation generation process. These changes need to be reviewed and validated so as to ensure the RefSeq database continues to be a reliable and high-quality gene annotation resource for the research community. Similarly, such review should be conducted for other gene annotation databases as well.

The research findings from this study also have an implication for the quantification of RNA-seq data generated by the recently emerged single-cell sequencing technologies.

Same as the quantification of bulk RNA-seq data, an accurate gene annotation is also required for quantifying single-cell RNA-seq data. It is therefore important to understand if and how the annotation choice impacts the quantification accuracy of the single-cell RNA-seq data as well.

