## Supplementary Figures S1-S5 for "Impact of gene annotation choice on the quantification of RNA-seq data"

This document includes Supplementary Figures S1-S5.

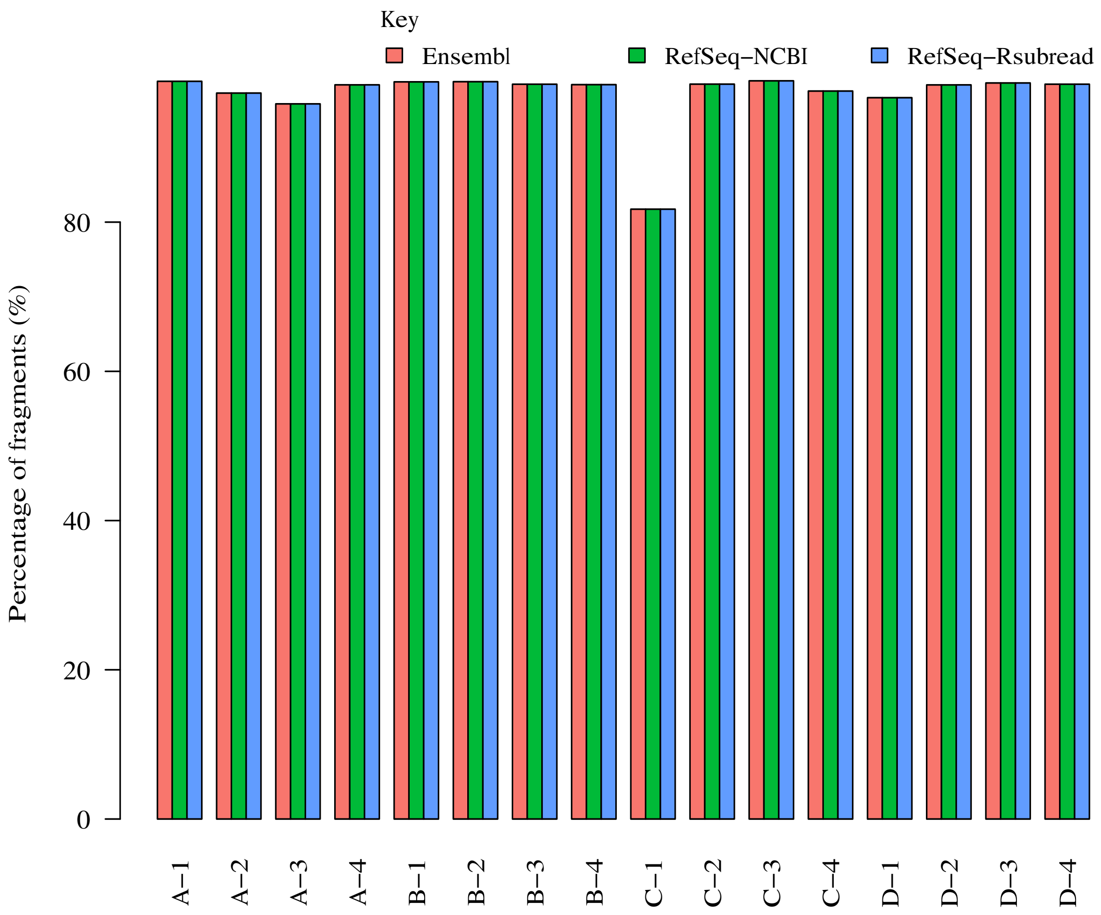


Figure S1. Percentage of fragments that were successfully aligned to the human reference genome GRCh38/hg38 in each library.

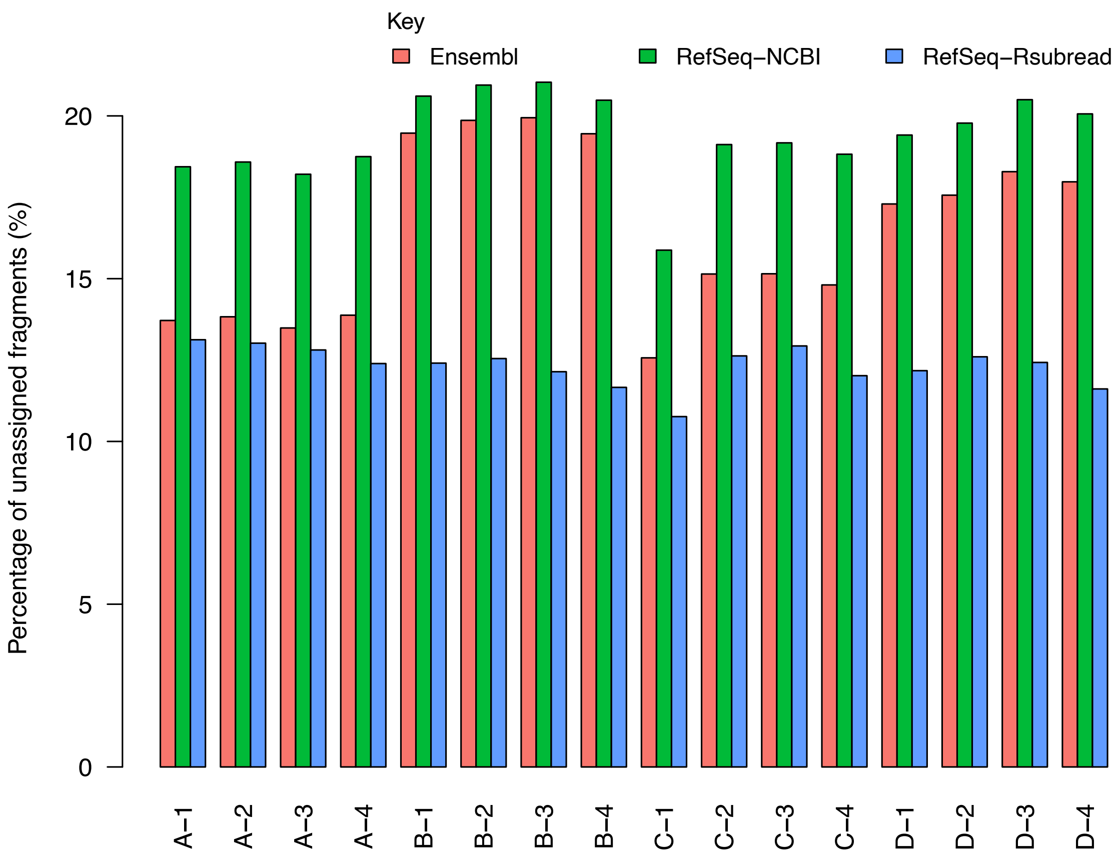


Figure S2. Percentages of fragments that failed to be assigned because they did not hit any annotated exons in an annotation. The percentage is calculated as the number of unassigned fragments divided by all the fragments included in a library.

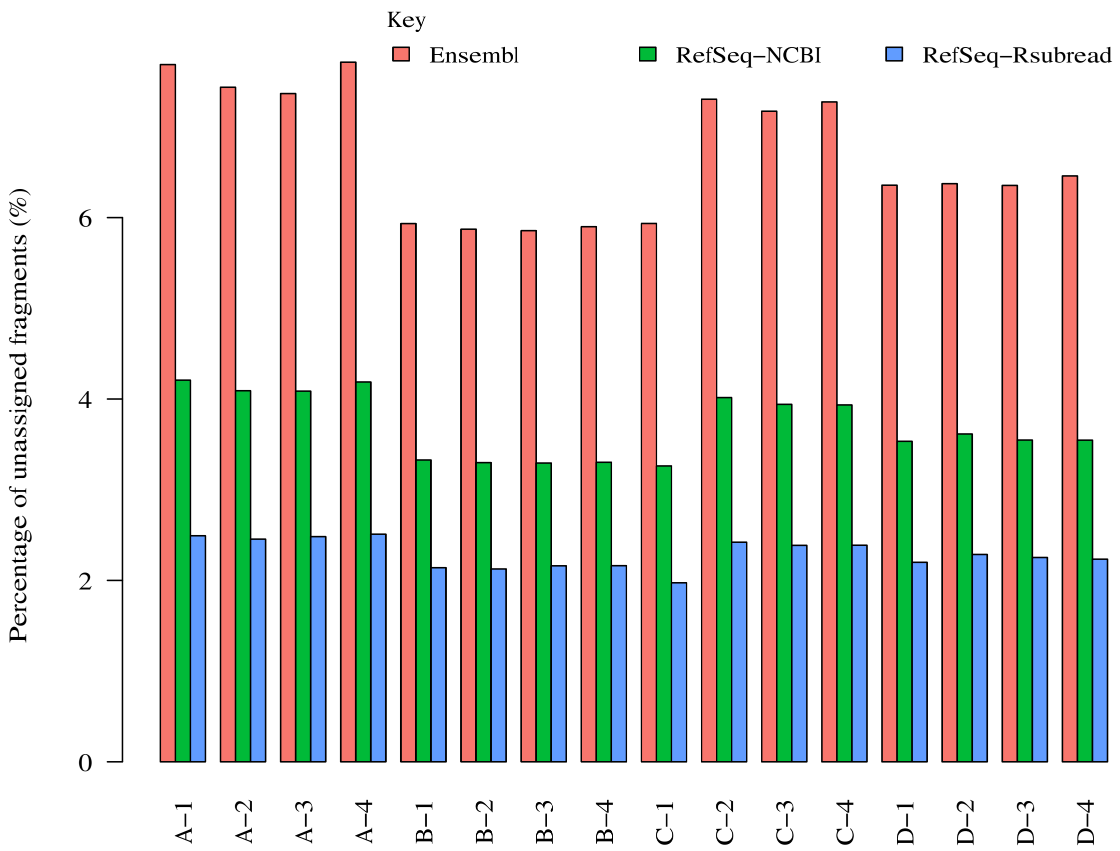


Figure S3. Percentages of fragments that failed to be assigned because of overlapping more than one gene in an annotation. The percentage is calculated as the number of unassigned fragments divided by all the fragments included in a library.

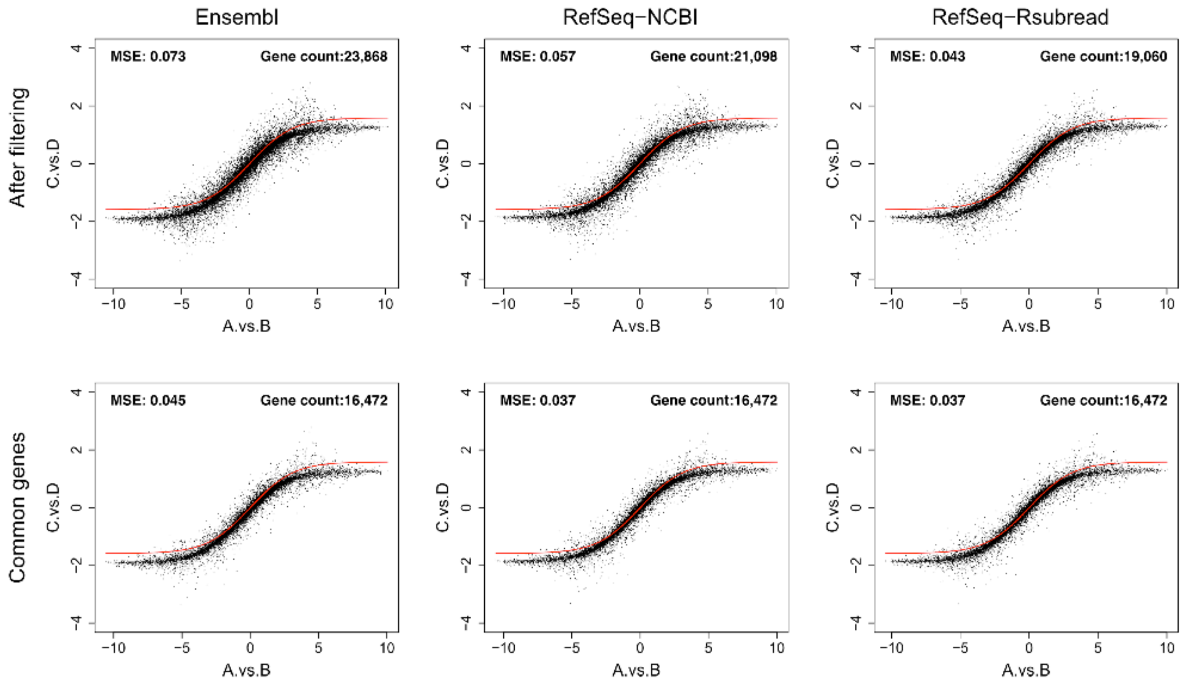


Figure S4. Titration monotonicity plots generated from using TMM normalized data. The red curve in each plot represents the reference titration. The Mean Squared Error (MSE) between the reference titration and the actual titration was calculated for each annotation, using all the genes that remained after filtering for lowly expressed genes (top row) or using common genes between the three annotations after filtering for lowly expressed genes (bottom row). In each plot, the horizontal axis represents the $\log_2$ fold changes of gene expression between sample A and sample B and the vertical axis represents the $\log_2$ fold changes of gene expression between sample C and sample D.

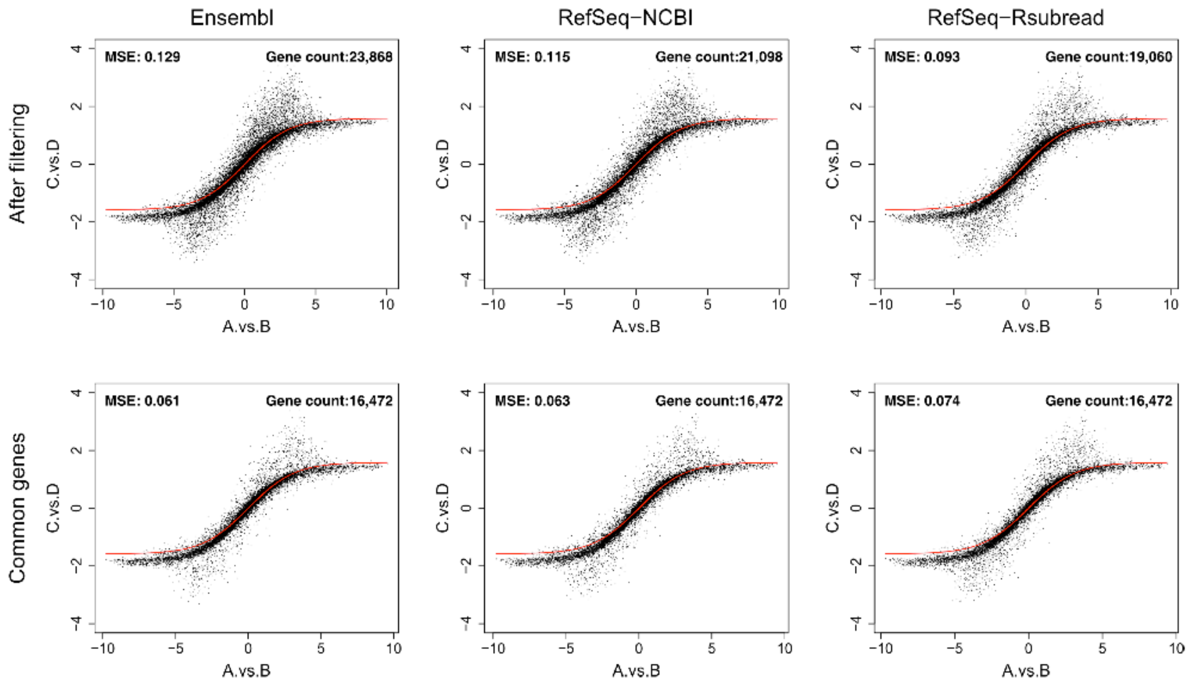


Figure S5. Titration monotonicity plots generated from using quantile normalized data. The red curve in each plot represents the reference titration. The Mean Squared Error (MSE) between the reference titration and the actual titration was calculated for each annotation, using all the genes that remained after filtering for lowly expressed genes (top row) or using common genes between the three annotations after filtering for lowly expressed genes (bottom row). In each plot, the horizontal axis represents the $\log_2$ fold changes of gene expression between sample A and sample B and the vertical axis represents the $\log_2$ fold changes of gene expression between sample C and sample D.
